# Choice of Parcellation Atlas Might Not be Too Critical for Connectomic Analysis

**DOI:** 10.1101/2022.12.20.521276

**Authors:** Steven C. Nesbit, Drew Parker, Ragini Verma, Yusuf Osmanlıoğlu

## Abstract

Connectomics has been a rapidly growing discipline in neuroimaging and neuroscience that evolved our understanding of the brain. Connectomics involves representing the brain as a network of regions, where the parcellation of the brain into regions using a template atlas is an integral part of the analysis. Over developmental and young adult cohorts of healthy individuals, we investigated how choosing parcellation atlases at certain resolutions affect sex classification and age prediction tasks performed using deep learning on structural connectomes. Datasets were processed on a total of 35 parcellations, where the only significant difference was observed for age prediction on the developmental cohort with a slight improvement on higher resolutions. This indicates that choice of parcellation scheme is generally not critical for deep learningbased age prediction and sex classification. Therefore, results between studies using different parcellation schemes could be comparable and repeating analyses on multiple atlases might be unnecessary.

## 1. INTRODUCTION

Connectomics is the analysis of the brain as a network of structurally and functionally interconnected brain regions. Within recent times, it has proven useful in understanding the healthy brain as well as various brain related diseases and disorders including autism, Alzheimer’s, and traumatic brain injury [1]. Connectomes are typically represented as graphs with brain regions as nodes and either structural or functional connectivity denoted as weighted edges. Structural connectivity is calculated over diffusion weighted MR images through a method called tractography, which models neural pathways connecting region pairs in the brain, while functional connectivity is calculated over functional MR images as the correlation of Blood Oxygenation Level Dependent (BOLD) signals among region pairs, denoting the pairwise activation of brain regions. Identifying regions of the brain, known as parcellation, is a necessary step in connectomic analysis to reveal graph nodes. Brain parcellation is commonly achieved by applying a template brain atlas over the brain image of individuals. A wide repository of parcellation atlases are available in the literature that differ on the type of features that the parcellation is obtained from such as functional or anatomical features [2]. Therefore, the choice of the atlas is an important question that the researcher needs to address before starting the analysis of the data.

A second problem that follows is the resolution of the atlas, that is, the number of regions into which the brain is parcellated. While some of the template atlases come in one resolution [3], others come in multiple resolutions [4]. In the face of this variation, it is necessary to evaluate how the choice of atlas and its resolution affects certain connectomic analysis. Previous research has sought to explore the effects of atlas choice on various graph-theoretical measures such as clustering coefficient, characteristic path length, global efficiency, and degree distribution [2, 5, 6]. Domhof et al. [2] also explored the effects of parcellation on phase oscillator and neural mass modeling and noticed significant heterogeneity in results between different atlases. However, the question remains as to how significant these differences are when it comes to certain connectomic analysis. In this paper, we address the problem of choice of atlas and resolution for two tasks, namely age prediction and sex classification, on two datasets over a wide range of atlas resolutions.

## 2. METHODS

### 2.1. Dataset and Preprocessing

We used diffusion wieghted images obtained from Human Connectome Project (HCP) [7] and Philadelphia Neurodevelopmental Cohort (PNC) [8] for our experiments. For the HCP, we investigated 200 subjects (104 females) from the young adult dataset in age range [22,36]. Structural connectomes were obtained from two different processing pipelines provided in EBRAINS [2] and braingraph.org [9]. The first pipeline on HCP [2] generated structural connectomes by using probabilistic tracking with 10M streamlines, over 20 different parcellation atlases that vary in regions of interest (ROI) ranging 31 to 294. Adjacency matrices were then created using the regions specified by each atlas as nodes and fiber counts as the edge weights. The second pipeline on HCP generated structural connectomes using probabilistic tracking on 1M streamlines, and repeated the process ten times to consider their average as the final connectomes. Connectomes were parcellated into five different resolutions of the Lausanne atlas [10] having 83 to 1,015 nodes. The reader is referred to [2] and [9] for further details on the processing pipeline of the connectomes.

The PNC is a developmental dataset consisting of 987 healthy individuals (546 females) in age range [8,22]. Structural connectomes derived from diffusion MRI data of PNC dataset were generated using probabilistic tracking [11] with 10M streamlines at ten different resolutions of the Schaefer atlas [4] having 120 to 1,020 nodes. Weights were then calculated for each streamline using SIFT2 [12] and connectivity between regions of interest was determined as the total number of streamlines connecting them. Adjacency matrices were created with brain regions as nodes and fiber counts connecting them as edge weights.

### 2.2. Deep Learning

The neural network consisted of three fully-connected layers of shapes that allowed for the maximum number of parameters to still fit the model, given the size of the datasets, with the matrices of the greatest number of nodes, in 8 GB of GPU RAM (Figure 1). It consisted of an input layer for a (*D, D*) matrix of size (*D*^2^,256), followed by a ReLU activation function. The next layer was of size (256,64), which was also followed by a ReLU activation function. The output layer was of size (64,2) for sex classification and (64,1) for age prediction. Each layer was followed by a dropout layer with 20% probability for the sex classification task, while only the first layer was followed by a dropout layer with 10% probability for the age prediction task. Overfitting was not observed after adding these dropout layers.

**Fig. 1:**
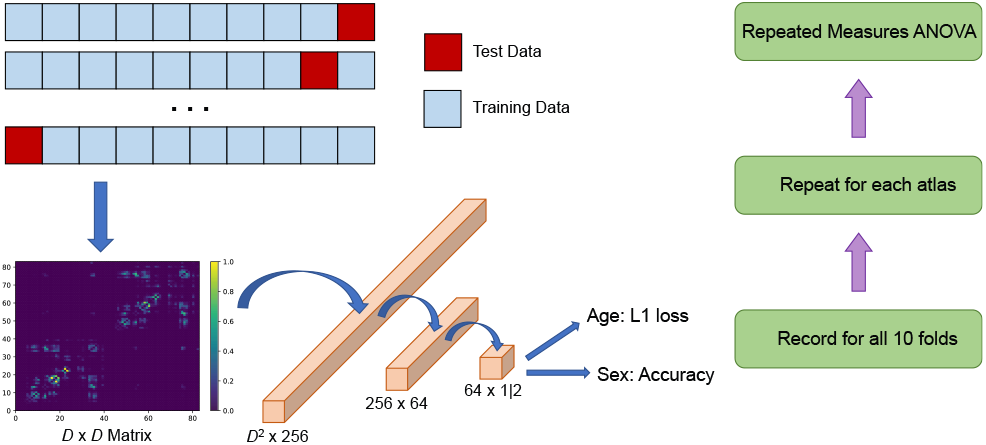
Overall summary ofthe experiments and the structure ofthe deep learning model.

To avoid connectivity bias across ages and sexes, connectomes, which are simply weighted adjacency matrices, were normalized by dividing each matrix element by the mean value of all elements for that matrix. For sex classification, each element was then divided by the maximum value of that matrix so that the input values to the neural network would be between 0 and 1. To meet the memory requirement for the largest adjacency matrices, mini batches of size 8 were used for all training regimens. Ten-folds cross validation was performed to determine sex classification and age prediction accuracies across the whole datasets. The neural networks were trained for 150 epochs using Adam optimization. Training for the sex classification task used cross-entropy loss, while training for age prediction used a mean squared error objective function. Sex classification performance was evaluated as percentage correct, whereas age prediction performance was evaluated using an L1 loss function.

### 2.3. Statistical Analysis

Repeated measures analysis of variance (ANOVA) tests were performed to determine if there were any significant main effects of the parcellation scheme choice on the sex classification or age prediction accuracy on the PNC and HCP datasets. Repeated measures ANOVA was used because the subjects in each fold were identical across atlases for a given dataset. Mauchly’s tests of sphericity were conducted to ensure the equality of variance assumption was not violated. In the case of age prediction error for the PNC dataset, equality of variance was violated as indicated by a Mauchly’s test of sphericity *p* < 0.05. Due to ϵ < 0.75, a Huynh-Feldt correction was applied to the ANOVA. Post-hoc tests consisted of dependent t-tests with Bonferroni correction.

## 3. RESULTS

### 3.1. Age Prediction

The top plot of Figure 2 shows the age prediction error, in years, for each atlas of the HCP dataset. The connectomes from the EBRAINS dataset are shown in black and braingraph.org in yellow. Mean age prediction errors, as shown by the red bars in Figure 2, ranged from 2.59 to 2.79 years. The atlas that achieved the minimum age prediction error was the Lausanne atlas with 129 nodes, whereas the atlas that produced the maximum error was the Harvard-Oxford 96 atlas. However, a one-way repeated measures ANOVA showed no significant main effect of connectome atlas selection on age prediction error for the HCP subjects (*F*(9,240) = 1.52, *p* = .063, *η*^2^ = .15). The top part of Figure 3 shows the age prediction error, in years, for each atlas of the PNC dataset, indicated in blue, with asterisks denoting significant differences. As shown by the red bars, mean age prediction errors ranged from 1.43 to 1.55 years. The Schaefer atlas that achieved the lowest age prediction error was the one with 820 nodes and the one that produced the highest age prediction error was the one with 120 nodes. The variances of the age prediction error for the different atlases were not equal, which was verified by Mauchly’s test of sphericity (*W* = .000, *ϵ* = .537, *p* < .001), so a Huynh-Feldt correction was made. A one-way repeated measures ANOVA with Huynh-Feldt correction showed a large and significant main effect of atlas on age prediction error (*F*(5,90) = 5.31, *p* < .001, *η*^2^ = .37). Post-hoc dependent t-tests with Bonferroni correction showed the Schaefer atlas with 120 nodes gave significantly higher error (*M* = 1.55, *SD* = 0.20 years) than the Schaefer atlas with 920 nodes (*M* = 1.43, SD = 0.14 years, *p* = .024) and the one with 1,020 nodes (*M* = 1.43, *SD* = 0.17 years, *p* = .005).

**Fig. 2:**
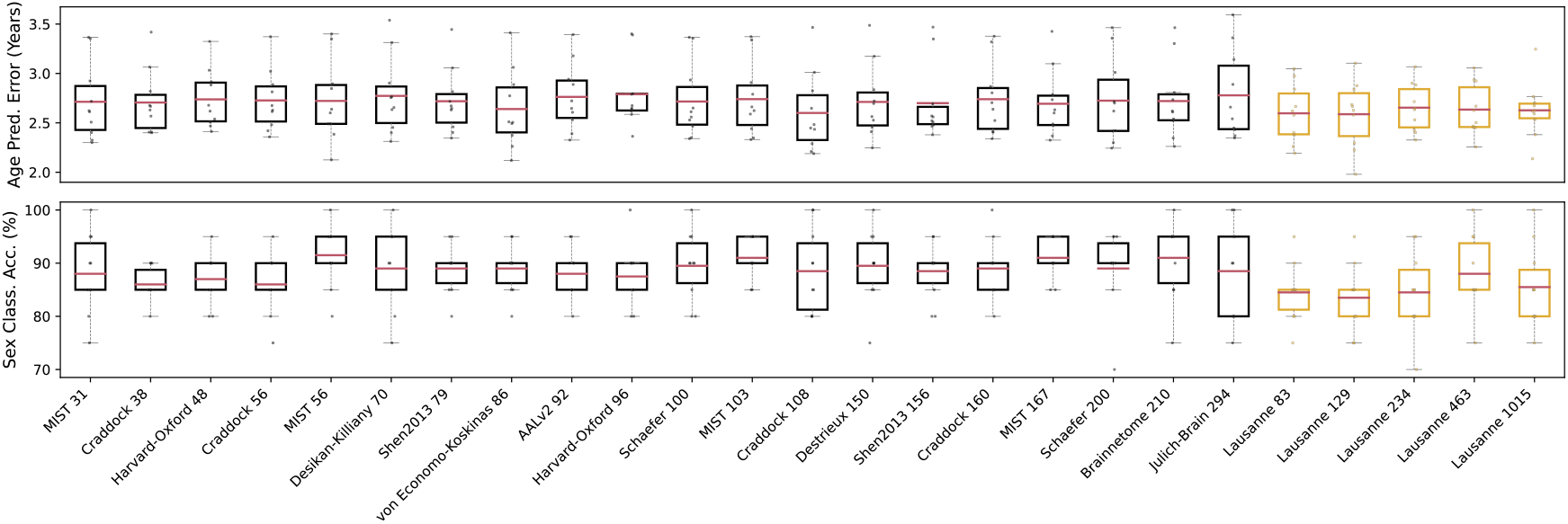
Age prediction (top) and sex classification (bottom) results on the HCP dataset. There were no significant differences observed for this dataset.

**Fig. 3:**
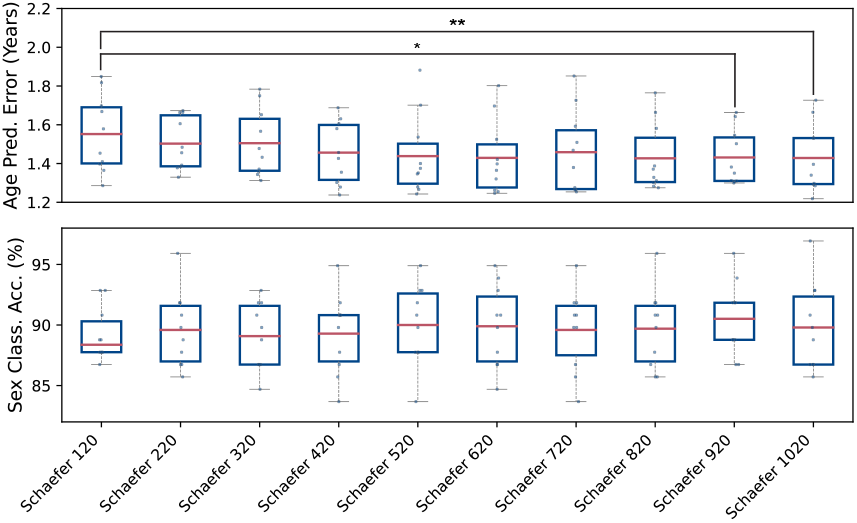
Age prediction (top) and sex classification (bottom) results on the PNC dataset. * indicates significance at *p* < .05 and ** indicates significance at *p* < .01.

### 3.2. Sex Classification

The bottom plot in Figure 2 shows sex classification percent accuracy for each atlas of the HCP dataset. Mean sex classification accuracies ranged from 83.5 to 91.5%. The atlas that achieved the best sex classification accuracy was the MIST 56 atlas and the atlas that produced the worst sex classification accuracy was the Lausanne atlas with 129 nodes. However, a one-way repeated measures ANOVA showed no significant main effects of connectome atlas on sex classification accuracy for the HCP subjects (*F*(9,240) = 1.52, *p* = .062, *η*^2^ = .15). The bottom half of Figure 3 shows the sex classification percent accuracy for each atlas of the PNC dataset. Mean sex classification accuracies ranged from 88.4 to 90.5%. The Schaefer atlas that achieved the best sex classification accuracy was the one with 920 nodes and the one that produced the lowest sex classification accuracy was the one with 120 nodes. However, a one-way repeated measures ANOVA showed no significant main effect of atlas on sex classification accuracy (*F*(9, 90) = 1.99, *p* = .051, *η*^2^ = .18).

## 4. DISCUSSION AND CONCLUSION

The only significant differences detected in this study were between the Schaefer atlas with 120 nodes and the Schaefer atlases with 920 and 1,020 nodes for age prediction error on the PNC dataset. However, the differences in age prediction error between connectomes parcellated with these atlases were no more than 0.12 years, indicating satisfactory performance for the Schaefer 120 atlas, despite its significantly higher error. Therefore, it is possible that parcellation atlas selection is not critical for deep learning tasks involving age prediction or sex classification of healthy subject connectomes. In addition, this indicates that it may be possible to compare works between studies that used different parcellation regimes.

Whereas sex classification accuracies were similar between the HCP and PNC datasets, the age prediction error was lower for the PNC dataset than it was for the HCP dataset. This may be due to the differences in age range for the two datasets, with the HCP subset containing participants aged 22 to 36 years, and the PNC dataset containing participants aged 8 to 23 years. The discrepancy in error may be caused by more significant structural changes in the brain present during development than during adulthood [13].

Although we showed that choice of parcellation scheme may not matter for deep learning sex classification and age prediction tasks, further research is warranted to explore tasks such as clustering of individuals into subgroups by using connectomes. The study also needs to be expanded over individuals with brain diseases or disorders as our analysis evaluated only healthy individuals. Additionally, we only considered choice of parcellation over structural connectomes. Thus, future work will incorporate functional connectomes as well in order to determine how well different parcellation schemes represent functional connections.

For sex classification and age prediction deep learning tasks over structural connectomes, the choice of atlas generally does not significantly affect performance. Therefore, when performing these tasks, the researcher may be better suited considering parcellation resolutions that do not include an unnecessarily large number of nodes. Since memory, storage, and computation time requirements for the connectomic analysis scale quadratically with the number of nodes in the parcellation, using a high resolution parcellation may delay connectomics research without producing much tangible benefit. In addition, it may not be necessary for a researcher to repeat analyses using multiple atlases, as the results here suggest that any reputable atlas should be able to produce robust results.

## 5. COMPLIANCE WITH ETHICAL STANDARDS

This research study was conducted retrospectively using human subject data made available in open-access by the Human Connectome Project (HCP) [7]. Ethical approval was not required as confirmed by the license attached with the open access data.

## 6. ACKNOWLEDGMENTS

Data were provided by the Human Connectome Project, WU-Minn Consortium (Principal Investigators: David Van Essen and Kamil Ugurbil; 1U54MH091657) funded by the 16 NIH Institutes and Centers that support the NIH Blueprint for Neuroscience Research; and by the McDonnell Center for Systems Neuroscience at Washington University.

No funding was received for conducting this study. The authors have no relevant financial or non-financial interests to disclose.

